# *RePAIR*: a power solution to animal experimentation

**DOI:** 10.1101/864652

**Authors:** V. Bonapersona, H. Hoijtink, RELACS, R.A. Sarabdjitsingh, M. Joëls

## Abstract

Low statistical power challenges the reliability of animal research; yet, increasing sample sizes to the required level raises important ethical and practical issues. We present an alternative solution, *RePAIR*, which capitalizes on the observation that control groups in general are expected to be similar to each other. As shown in a simulation study, including information of previous control experiments in the statistical analysis using *RePAIR* reduced the required sample size by 49% or increased power up to 100%. We validated the potential of *RePAIR* in a unique dataset based on 7 independent experiments across the world, studying cognitive effects of early life adversity in mice. *RePAIR* comes with an open-source web-based tool and can be widely used to largely improve quality of animal experimentation.

**One Sentence Summary:** Prior studies’ information can reduce use of animals or increase statistical power, improving animal research reliability

## Main Text

Researchers embarking on a new experimental study calculate in advance the number of animals (sample size) required to allow distinction between a *real* effect and a chance finding. This calculation depends on statistical power (“Key concepts” in Suppl Methods), which is frequently set *a priori* at 80%; thus, it is expected that 8 out of 10 studies investigating a real effect will correctly conclude that the effect exists (true positive), while 2 will not (false negative). As power decreases, the rate of false negative as well as false positive results will increase(*1*). Prospective study power is therefore directly related to the reliability of research.

The prospective power assumption of 80% has been seriously challenged. In a landmark paper from 2011 based on 48 meta-analyses across Neurosciences, Button and colleagues showed that the median power was estimated at 20%, rather than 80%(*2*), in agreement with previous reports in psychology(*3*). We confirmed and extended this finding to a much larger sample of preclinical studies(Fig.1-A), both in the field of Neuroscience and Metabolism(Fig.S2). This power problem contributes significantly to the reproducibility crisis in preclinical research(*4*): single, underpowered studies have a low chance to detect a real effect(*1*), although when analyzed meta-analytically they can still be informative(*5,6*).

**Fig. 1.**
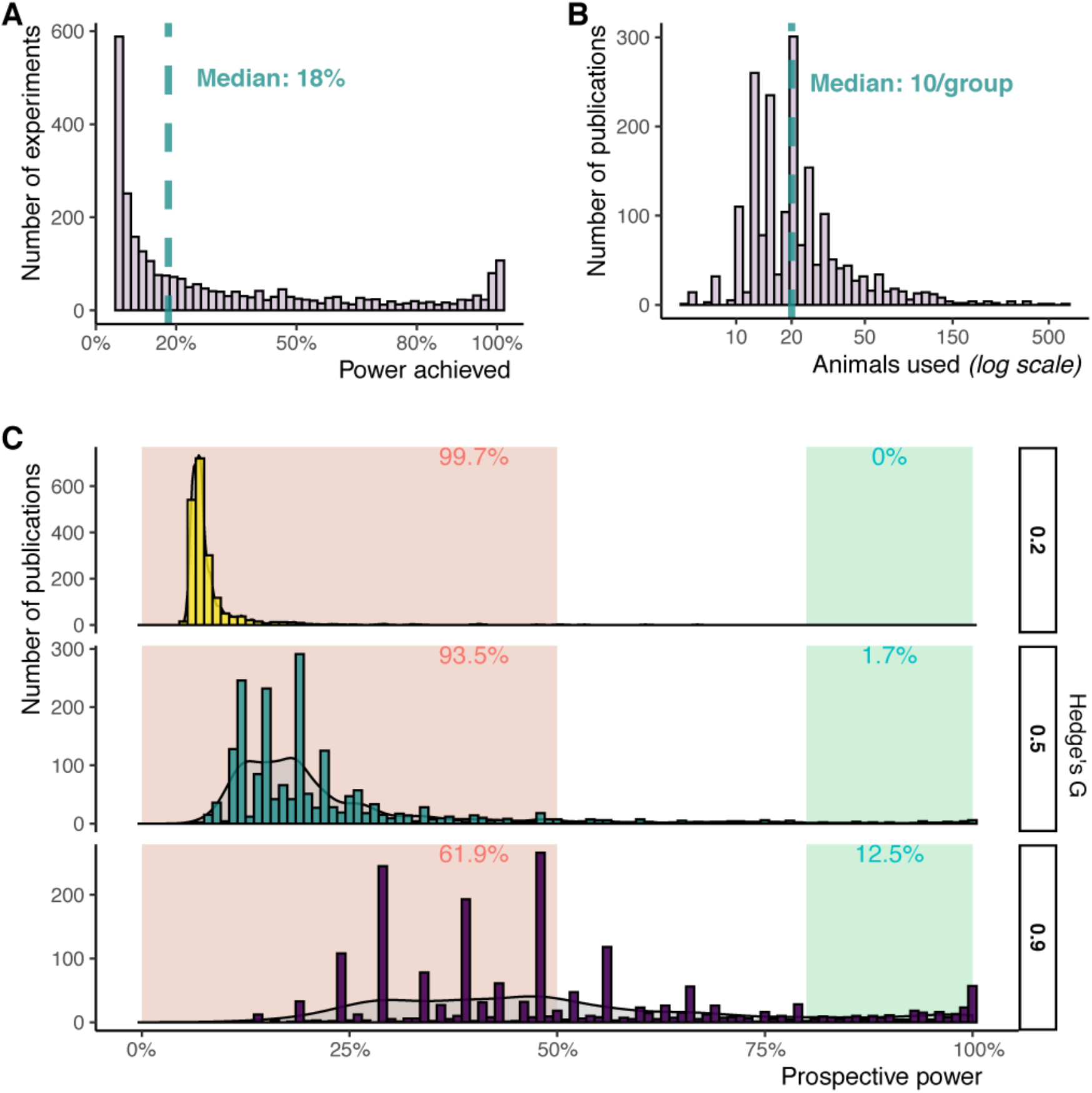
Preclinical experiments are severely underpowered. Data was collected by conducting an EMBASE comprehensive literature search on April 12^th^ 2019 that identified 1935 primary publications in mice and rats used in 69 meta-analyses (Suppl. Methods). (**A**) Histogram of power achieved by identified experiments (Welch t-test, effect sizes from publications, Fig.S1”DataB”). Green: median equal to 18%. (**B**) Histogram of animals per study when considering the two largest independent groups (Fig.S1”DataA+B”). Green: median equal to 20 (i.e.~10 animals per group). (**C**) Prospective power of studies when considering a range of common effect sizes (Fig.S3, Fig.S1”DataB”) and assuming at least one sufficiently powered experiment per publication. The highest peaks in the histograms are due to a non-uniform distribution of animals used as shown by Fig.1-B. Histogram and density plot of the same data are overlapping. In red: power ≤50%; in green: power ≤80%.

Why is the actual study power so much lower than assumed *a priori*? One explanation is that effect sizes are often estimated optimistically, based on earlier findings that are liable to (publication) bias(*7*). A second explanation is that rodent experiments are frequently exploratory in nature(*8*), and many scientists opt to use a debatable “standard” 6 to 10 animals per group. When considering a Welch independent samples t-test (alpha=0.05) and 10 animals per group (Fig.1-B), one would need to assume an effect size *Hedge’sG*=1.4 to reach a power of 80%. Such an expected effect size is far larger than what is commonly observed in rodent literature(Fig.S3).

**Fig. 2.**
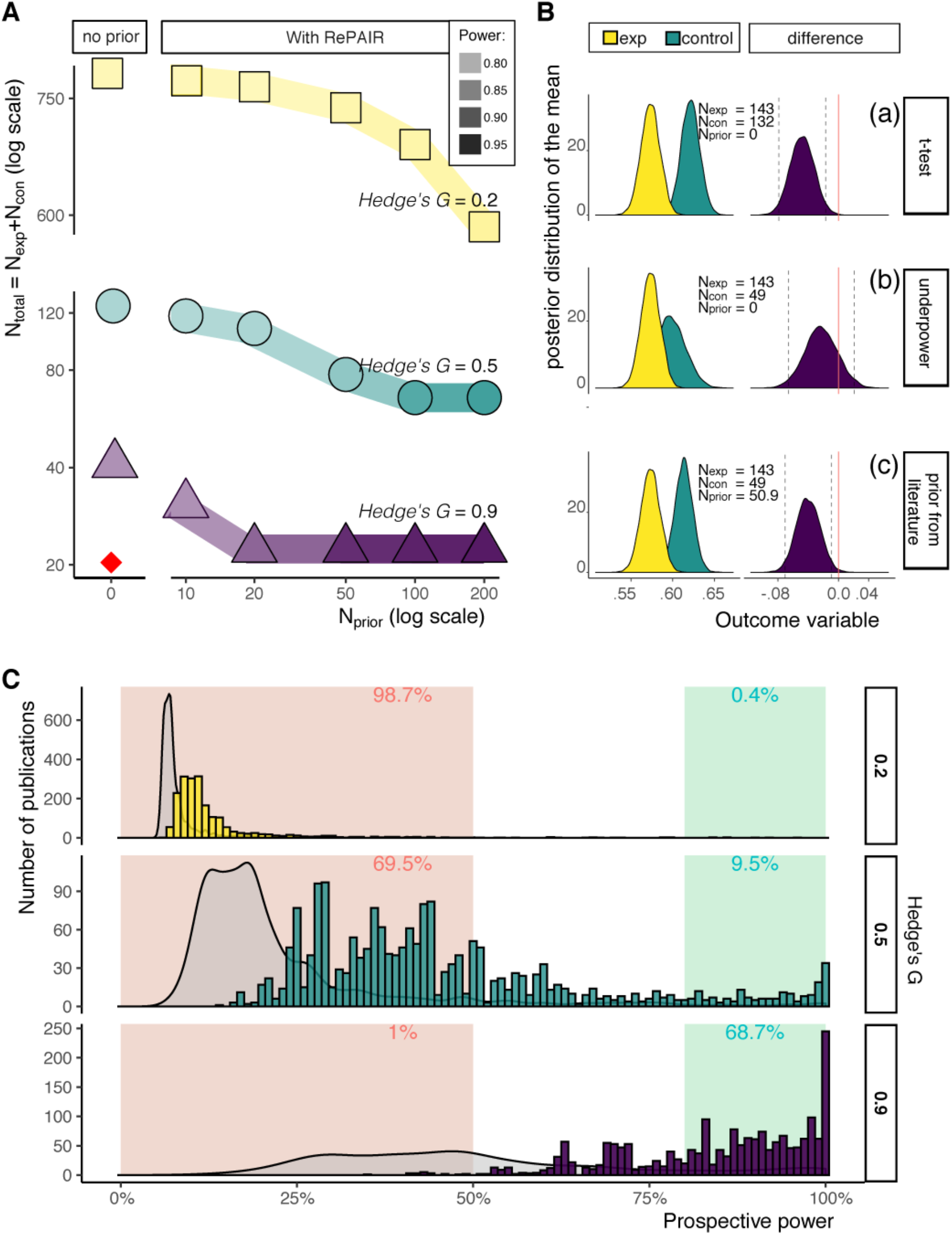
RePAIR can decrease the number of animals required for sufficiently powered research. (**A**)Simulation study on the relationship between prior (*index*=1), sample size and power. A n_prior_ equal to 0 corresponds to a standard sample size estimation (Welch t-test: two independent samples, alpha=0.05, effect sizes as in Fig.1-C, power=80%). The red diamond indicates the current median sample size. An increase in color intensity signifies an increase in power. As n_prior_ increases, n_total_ decreases until a plateau is reached. Subsequent increases in n_prior_ will result in increased prospective power. (**B**)Application of RePAIR to the experimental dataset RELACS. Posterior distributions of each group and of their mean difference. The test is significant if 0 (red line) is outside the 95% confidence interval (dashed lines) of the means’ difference distribution. From the top (Table S4), **(a)**RePAIR analysis without prior provides the same result as a Welch t-test; **(b)**if ^n^con is decreased, the study becomes underpowered; **(c)**but this can be rescued if a prior from (unrelated) published literature is introduced. (**C**)Prospective power when using RePAIR with an *index=0.3* (only 30% of prior information is used, i.e. n_prior_=0.3*n_con_ of other studies within the same meta-analysis (Fig.S1,”DataA”)) but maintaining current resources (n_total_ kept the same; n_con_=n_total_/3 as recommended rule of thumb;) shown as histogram. Grey density plots represent the current prospective power as Fig.1-C.

Effect sizes should be estimated specifically for each research question, but what is a plausible range that a scientist could expect? A sensible approach is to consider the frequency of effect sizes generally observed. We considered the distribution of 2738 effect sizes retrieved from 482 primary articles (Fig.S1,”DataB”); this was replicated in a separate dataset(*2*). We defined the quantiles of 0.25, 0.5 and 0.75 as small, medium, and large effect sizes respectively. These corresponded to *Hedge’sG* of 0.2, 0.5, 0.9(Fig.S3), which is almost identical to *Cohen’s d* rule of thumb for small, medium and large effect sizes(*9*). Some subfields in preclinical research may be placed more towards the lower (e.g. behavioral research(*5*)) or higher end of this distribution (e.g. molecular studies(*10*)). Nonetheless, this range can be used to estimate the prospective power that a scientist currently starting a new project could expect.

Considering the estimated *Hedge’sG* values for small, medium, and large effect sizes respectively, which percentage of rodent studies are then -prospectively- sufficiently powered? For this, we selected data of 1935 rodent primary studies derived from 69 meta-analyses, identified with a systematic search, independently of research field(Fig.S1,”DataA”). We extracted the sample size only of the two largest groups reported in each paper, assuming that at least the comparison of these two groups were sufficiently powered while all other experiments may have been control experiments. This yields a best-case scenario, and prior to any potential subsequent multiple testing, experimental bias, p-hacking/fishing, selective reporting, etc. Even in this best-case scenario, only 12.5% of this large sample of rodent studies were sufficiently powered(Fig.1-C, large effect size).

This power problem could be solved by increasing the number of animals per experiment. For example, the effect sizes of *Hedge’sG* 0.2 and 0.9 would correspond to a requirement of 394 and 21 animals per group respectively, when considering an independent samples t-test (alpha=0.05, power=80%). This clearly challenges the societal debate on reducing the number of animals to conduct scientific research, quite apart from practical and financial considerations. How can sufficient power be reached without increasing the number of animals per group to unrealistically high levels?

To address this dilemma, we here propose a novel approach in preclinical research, which capitalizes on the observation that new studies are rarely disconnected from earlier ones using the same experimental endpoints. In research practice, any new experiment is planned based on information from previous studies – either from literature or one’s own. Previous knowledge can be incorporated within statistical analysis by using Bayesian priors, namely distributions that describe the mean and variance of an experimental outcome.

Despite the advantageous properties of priors such as in sample size calculations(*11*), their use is often criticized and dismissed because priors represent a naturally subjective state of knowledge rather than objective principles(*12*). However, compared to human studies, rodent experiments have the great advantage to be relatively well-controlled and to use standardized tests. Even though variations between strains and labs definitely exist(*13, 14*), researchers have similar expectations about how a control group “should respond”. This prior knowledge is so important that a researcher would not trust his/her own or other’s data and judge that an experiment “did not work” or “needs to be better optimized” if the expectation is not met. If this common research practice is translated into statistical terms, one would state that researchers assume control animals always to belong to the same population. We formalized this property and developed *RePAIR (Reduction by Prior Animal Informed Research)*, a statistical method that uses previously obtained information to limit the number of animals, yet perform well-powered research.

The acceptance of this assumption in daily research practice warrants the formal statistical use of priors to supplement the data of the control group. Following this line of reasoning, we developed an algorithm (“Theory”, Suppl. Methods) to address the dilemma sketched above, based on the formula:

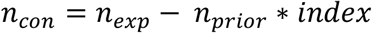

where the number of animals of the control group (*n_con_*) can be reduced by the number of control animals from prior studies (*n_prior_*) multiplied by a weight (*index*, value between 0 and 1) that describes the similarity between control and prior groups, while the experimental group (*n_exp_*) remains the same. It follows that the number of animals necessary in the control group is effectively diminished by the weighed prior.

With this formula, we performed a simulation study to estimate how the use of priors relates to sample size and power (Fig.2-A). Although it is theoretically possible to decrease *n_con_* to 2 for standard deviation calculation, this is not advisable as it would challenge experimental blinding and randomization. As a rule of thumb, we recommend that at least one third of the total number of animals consists of controls. Nonetheless, a further increase in *n_prior_* can be beneficial as power could be enhanced up to its highest boundary, i.e. 100% with large effect sizes.

In the simulation, the *index* was set equal to 1, meaning that prior and control animals were assumed to be interchangeable. By changing this index, scientists themselves (by *expert elicitation*, an accepted practice in Bayesian statistics(*15*)) can decide to what extent they value earlier data; importantly, even conservative (low) indices will be beneficial. By selecting earlier data, scientists define *prior distributions*, meaning summaries of previous studies with respect to the mean of the control group. The selection of previous experiments and related indexes may be criticized because it introduces subjectivity in the statistical analysis. In other words, the prior distribution may not be a good description of the population. To address this concern, we performed a sensitivity simulation study (“Sensitivity simulation”, Suppl. Methods), where we verified that variations in prior specifications due to random sampling consistently provide better prospective power than current practice (Fig.S4). Despite its apparent robustness, researchers should be mindful when specifying the prior. Yet, as the number, quality and consistency of experiments increase, so will the quality of the prior.

We next tested the validity of this method for one particular experimental endpoint – i.e. the effects of early life adversity (ELA,(*16*)) on spatial learning in adult male mice. Due to the lack of sufficient power of single studies, the experimental dataset was gathered by aggregating experiments (that in principle shared the same design) from several laboratories around the world – the *RELACS (Rodent Early Life Adversity Consortium on Stress)* consortium. Information of over 250 animals was collected, which was required to verify a significant effect of ELA on memory (*t(3)=272.99, p=0.003*) according to our prospective power calculation (“RELACS dataset”, Suppl. Methods). To estimate the control prior, a researcher (VB) randomly selected relevant yet unrelated (non-ELA) publications on spatial learning using this particular experimental task, blinded to their results (“Prior specification”, Suppl. Methods). By applying this prior from literature, we reached the same conclusion as derived from the aggregated experimental dataset, but now with 49 instead of 132 control animals (Fig.2-B). In other words, this same experiment could be conducted with 83 fewer animals, while maintaining a power >80%.

To facilitate the use of the algorithm, we created an open-source web-based tool (https://vbonapersona.shinyapps.io/repair_app_submit/) that allows anyone designing future experiments to improve the quality of his/her studies. With a user-friendly interface, one can 1) calculate (multiple) prior parameters from summary statistics, 2) perform sample size calculations and 3) execute analyses. R scripts are also publicly available (https://osf.io/wvs7m/).

Besides the theoretical value shown by the simulation study (Fig.2-A), the robustness from the sensitivity simulation (Fig.S4), and the applicability demonstrated on a real-life dataset (Fig.2-B), the immediate potential impact of this approach can be evaluated by applying it to metrics of current practice. Despite the difficulty of estimating how much prior knowledge exists in the literature, it can be assumed that if publications are similar enough to be included in a metaanalysis, they are also similar enough to be used for priors. We recalculated the prospective power of Figure 1-B. Controls of other studies within the same meta-analysis were used as prior, while new experiments were simulated with resources equal to the ones currently available (*n_total_* kept equal) re-distributed in favor of the experimental group (Fig.2-C). For *Hedge’sG=0.9*, application of *RePAIR* improved the current 12.5% of sufficiently powered studies to 69%. These calculations were performed with an *index* of 0.3, although little would change if an *index* of 1 would be used instead (Fig.S5).

In summary, there is a growing awareness of reproducibility issues in animal experimentation which is only partially addressed by preregistration and the introduction of more rigorous guidelines (e.g.ARRIVE). We here introduce a statistical method, *RePAIR*, that uses previously obtained information in order to limit the number of animals, yet perform well-powered research or reach higher statistical power with the same total number of animals. Although here discussed in relation to t-tests, the same ploy can be extended to more complex experimental designs (e.g.2×2-ANOVAs) where multiple groups could then be considered as “controls”, or to human studies where the control group is sufficiently homogeneous. Nonetheless, this is clearly only applicable to experimental paradigms that are frequently used. Small exploratory studies will remain indispensable to open up new avenues and develop new techniques.

Bayesian priors have been criticized because of the subjectivity they introduce. We argued that especially in animal research one can assume that previously used subjects in essence belong to the same population, since interpretation of experimental results is often bound to previous expectations of the control group. For commonly used outcomes, priors can be openly discussed amongst panels of experts, an accepted approach in Bayesian statistics(*15*). We foresee a role for scientific societies in supporting this process, since they can not only provide an open source platform to collect previous datasets of commonly used tests in their field but also organize a critical mass of experts who jointly develop and maintain accepted priors for these tests. With the use of our web-based tool, these priors can then be used by everyone designing a new experiment, improving the overall quality of animal research. This will add to the value of animal experimentation in those cases where good alternatives are not available.

## Acknowledgments

We would like to thank Jelle Knop and Milou Sep for the helpful discussions, and Prof. Ron de Kloet for critically reviewing the manuscript.

## Author contributions

V.B. contributed with the conceptualization, data curation, analysis, investigation, methodology, software, visualization and writing the manuscript; H.H. contributed with the conceptualization, analysis, methodology, supervision, and reviewing/editing the manuscript; the RELACS consortium provided the data; R.A.S. contributed with the conceptualization, project administration, supervision and editing/reviewing the manuscript; M.J. contributed with the conceptualization, funding acquisition, project administration, supervision, and writing the manuscript.

## Data and materials availability

All data, code and materials used in the analysis are openly available and can be downloaded at https://osf.io/wvs7m/.

## Supplementary Materials

Materials and Methods

Figures S1-S5

Tables S1-S4

References (*17-22*)

