## Supplementary material for "*RePAIR*: a power solution to animal experimentation"

**This PDF file includes:**

Key concepts
Materials and Methods
Figs. S1 to S5
Tables S1 to S4
List contributors to the RELACS consortium

**Other Supplementary Materials for this manuscript include the following:**

Available at https://osf.io/wvs7m/:

Databases
R scripts

**Key concepts**

Sample size refers to the number of animals resulting from a power analysis executed with predefined characteristics to answer a research question. Animals in a group constitute a *sample* from a corresponding *population*. Statistical inference can be used to generalize findings from the sample to the population, provided that the sample is random and sufficiently large. In other words, experiments are performed to conclude something about the population(s), by gathering samples to estimate characteristics of the population(s).

Experimental design refers to how the animals are allocated to the different experimental conditions, which are defined by independent variables. Depending on the number of (in)dependent variables, the design can become more and more complex. For simplicity, we will consider throughout our study a simple and common experimental design: a comparison between a control and an experimental group. As typical (although questionable) in animal research, we will assume normality. The standard test in such circumstances is the Welch t-test, which is equivalent to the Student's t-test but does not assume homogeneity of variances.

The effect size quantifies the difference between two groups. In animal experiments with two independent groups, effect size is commonly measured as standardized mean difference, here calculated as *Hedge's G*, which is equivalent to *Cohen's d* (*17)*.

The alpha level defines the probability of having a "false positive" effect. It is normally set at 0.05, meaning that is expected that in 5 out of 100 experiments a researcher would wrongly conclude that there is an effect when in fact there is not.

The statistical power of a study defines the probability of detecting a "true positive" effect, and it is normally set at 80% (*18*). This means that if 100 experiments are performed, it is expected that in 80 of them a researcher would correctly conclude that an effect is present, and in 20 would wrongly conclude that there is no difference ("false negative").

A prior distribution summarizes information of previous studies with respect to the mean of the control group. A posterior distribution summarizes information of previous and current studies with respect to the mean of the control group. Of note, the mean and the variance of the experimental group also have a prior and a posterior distribution. However, the prior distribution is "uninformative", meaning that it will not have impact on the results. Therefore, the posterior distributions that describe mean and variance of the experimental group depend only on the information of the current experiment.

### Materials and Methods

#### Overview

Our methodology can be divided into four parts. Firstly, we describe the current scenario of animal research, with a specific focus on prospective study power (Section "Estimation of study power"). This was achieved by performing a systematic literature search, which yielded two datasets (Fig. S1). Throughout the manuscript, we specifically mention which dataset is used for which calculation.
Secondly, we developed *RePAIR*, which uses information of old experiments to limit the number of animals required in current experiment (Section "Using prior information"). Mathematical formulas and a simulation study are presented to show how *RePAIR* works.
Thirdly, we validated *RePAIR* (Section "Validation study") on a real-life dataset (RELACS, Sub-section "RELACS dataset"), which was achieved by the aggregation of (un)published data of multiple laboratories. Here, a prior from independent literature was used. Since prior specification may be criticized for its subjectivity, we performed a sensitivity simulation study to evaluate how variations in article selection may impact the results. Such variations were experimentally estimated with a sampling strategy.
Lastly, the applicability of this approach is investigated (Section "Immediate potential impact"). By using information from the systematic search, we simulated new experiments with the same resources as the current ones, and calculated their prospective power with *RePAIR*. The improvement in prospective power showcases the immediate potential impact of *RePAIR*.

#### Estimation of study power

Since *real* effect sizes are not known, estimating statistical power of animal research is equivocal. A common approach is to calculate *post-hoc* power (Sub-section "Post-hoc power") from meta-analyses identified with a systematic literature search (Sub-section "Data collection"). Although this approach replicated previous studies (*19*), it has major limitations (*20*). An alternative approach is to estimate a reasonable *prospective* power (Sub-section "Prospective power"), which aims to describe a plausible scenario for new experiments.

<u>Data collection</u>

To identify meta-analyses on rodent primary studies, a systematic literature search was conducted on April 12th 2019 in Embase. By screening titles, the search string *(rodent* OR mice OR mouse OR rat*) AND (meta-analys* OR metaanalys*)* identified 170 publications, while one additional record was identified via other sources. The articles' full-texts were screened by two authors (VB and RAS) and included if it matched the pre-defined inclusion criteria (Table S1). For a flow chart of the methodology, see Figure S1.

The identified meta-analytic articles ($n_{ma}$ = 69) were used to select primary publications in mice and rats. Of all included primary publications in each meta-analysis ($n_{primary\_study}$ = 1935, Fig. S1 "Data A"), we extracted the **sample sizes** of the two largest groups (equally split if only pooled quantities were reported), independent of the complexity of the experimental design, number of experiments and outcomes reported. We assumed that *at least* the comparison between these two groups would have been sufficiently powered.

Of the 69 identified meta-analyses, 8 meta-analyses matched our additional criteria for effect size estimation. These belonged to the fields of Neuroscience and Metabolism. From the resulting 482 primary studies, we extracted the summary statistics (mean, standard deviation or standard error of the mean, sample size) of all available comparisons between two independent groups, from which we calculated 2738 *Hedge's G* ($n_{summ\_stat}$, Fig. S1 "Data B").

Post-hoc power

From each set of summary statistics extracted, the achieved power was retrospectively calculated while assuming two independent groups (Welch t-test, two-tailed, alpha = 0.05, sample size and *Hedge's G* from Fig. S1 "Data B"). Post-hoc power is therefore the probability to reject the null-hypothesis (i.e., no difference between the control and experimental group) with the observed sample sizes if the effect size is set equal to the observed effect size.
This retrospective power calculation (Fig. 1A, Fig. S2) is a biased estimation of prospective study power, because it is a decreasing function of the p-value (*21*). Nonetheless, it is an impressive replicate of previous reports (*19, 22*), which used meta-analysis to estimate real effect sizes.

Prospective power

Besides the limitations of *post-hoc* calculations, we aimed to describe *prospective* power as a plausible range - rather than a single value - to mimic scenarios of researchers initiating a new study. The range of *prospective* power (Fig. 1-C) was calculated based on a range of effect sizes, while assuming two independent groups (Welch t-test, two-tailed, alpha = 0.05, sample size from Fig. S1 "Data A"). Prospective power is therefore the probability to reject the null-hypothesis with the observed sample sizes, if the effect size is set equal to a small, medium and large value, respectively.

To estimate a plausible range of effect sizes in preclinical literature, we calculated the 25$^{th}$, 50$^{th}$ and 75$^{th}$ percentiles of *Hedge's G*'s absolute values and defined them as small, medium and large effect sizes, respectively (Fig. S3, based on Fig. S1 "Data B"). Blinded to the results, we chose the 25-75% interval instead of the 95% confidence interval to avoid extreme values. Extremely low effect sizes may not be biologically relevant and are confounded by null effects, while extremely high values may lead to interpretation issues and are confounded by over-estimations due to biases. We confirmed (see R script at https://osf.io/wvs7m/) that these values are replicable by applying the same methodology to a separate dataset (*2, 10*).

#### Using prior information

Studying whether two numbers (A and B) are distinct is equivalent to investigating whether their difference (A – B) is different from 0. To calculate the difference distribution, we estimated the populations from the respective samples by using (un)informative priors. In sub-section "Theory", we describe how to compute a confidence interval for the difference in control and experimental means while using prior information from old experiments. In sub-section "Simulation study", we investigate how prior information can be used to decrease the number of animals required to answer a research question.

Theory
In the control group

$$y_i \sim N(\mu_{con},\ \sigma^2_{con}) \quad (1)$$

where $y_i$ denotes the score on the outcome variable in the control group for $i = 1, \ldots, n_{con}$ animals. Similarly, in the experimental group

$$y_i \sim N(\mu_{exp},\ \sigma^2_{exp}) \quad (2)$$

for $i = 1, \ldots, n_{exp}$. Dropping the subscripts $con$ and $exp$, the posterior distribution of $\mu$ and $\sigma^2$ in the control and experimental groups is given by (*section 3.2 and 3.3 of* (*21*))

$$g(\mu, \sigma^2) = g(\mu|\sigma^2)\ g(\sigma^2) = N(\mu|m_{post,}\ s^2_{post})\mathrm{X}^{-1}(n_{post}, s^2_{post}) \quad (3)$$

where the posterior mean is

$$m_{post} = \frac{n_{prior}}{n_{prior}+n} m_{prior} + \frac{n}{n_{prior}+n} \bar{y}\ , \quad (4)$$

the posterior variance is

$$s^2_{post} = (n_{prior} s^2_{post} + (n-1)s^2 + \frac{n_{prior} n}{n_{prior}+n} \left(\bar{y} - m_{prior}\right)^2 / n_{post}, \quad (5)$$

and the posterior degrees of freedom is

$$n_{post} = n_{prior} + n. \quad (6)$$

For the experimental group, the posterior distribution is based on an uninformative prior distribution, that is, the prior sample size $n_{prior} = 0$. For the control group, an informative prior distribution based on a previous study is used, where $n_{prior}$ denotes the sample size of the previous study, $m_{prior}$ denotes the mean of the scores on the outcome variable in the previous study and $s^2_{prior}$ the variance. In fact, the posterior distribution for the control group is based on $p = 1, \ldots, P$ prior studies. Bayesian updating is used to obtain the posterior distribution for the control group:

- **Step 1.** Use equation (3) to update an uninformative prior distribution ($n_{prior} = 0$) with the data from the first prior study $p = 1$ weighted with $n_{prior} = n_1 index_1$, where $n_1$ denotes the sample size of the first prior study, resulting in a posterior distribution.
- **Step 2.** This posterior distribution becomes the current prior distribution.
- **Step 3.** For $p = 2, \ldots, P$, that is, the remaining prior studies, use equation (3) to update the current prior distribution with the data from prior study $p$ weighted with $n_{prior} =$

$n_p index_p$, where $n_p$ denotes the sample size of the $p$th prior study, resulting in a posterior distribution that will have the role of prior distribution in the next step.

- **Step 4.** In this last step, using equation (3) the prior distribution resulting from Steps 1 through 3 is updated with all the data from the current study, rendering the posterior distribution in the control group.

The confidence interval for $\mu_{exp} - \mu_{con}$ is obtained by sampling $t = 1, \ldots, 10000$ values $\mu_{exp}^t$ and $\mu_{con}^t$ from the respective posterior distributions and computing their difference $\delta^t = \mu_{exp}^t - \mu_{con}^t$. The 2.5[th] and 97.5[th] percentile of the distribution of $\delta^t$ for $t = 1, \ldots, T$ constitute respectively the lower and upper bound of the 95% confidence interval for $\mu_{exp} - \mu_{con}$. When the value 0 is/is not contained in the confidence interval, the null-hypothesis that the experimental and control means are equal is not/is rejected.

Simulation study

We performed a simulation study (Fig. 2-A) to evaluate to which extent a prior could reduce the number of animals necessary and how this would influence study power. The more informative a prior for the mean in the control group, the more influence it will have on the conclusions of the experiment. Mean and variance of data in the control group were kept identical in all conditions ($\mu_{con} = 0, \sigma_{con}^2 = 1$); therefore, the influence of the prior was dependent only on its varying sample size $n_{prior}$. Table S2 summarizes all factors varied in the simulation. For each combination of factors, 10000 datasets were sampled from the corresponding population.

Firstly, we calculated how many animals ($n_{total} = n_{con} + n_{exp}$) one would need to perform experiments with the determined characteristics given a standard sample size calculation ($n_{prior}$ = 0), later confirmed by G*Power (*22*). This calculation assumed a balanced design, meaning $n_{con} = n_{exp}$. Secondly, we decreased $n_{con}$ by adding $n_{prior}$ while keeping $n_{exp}$ the same. Since it would be illogical for $n_{con}$ to become negative when $n_{prior} > n_{con}$, $n_{con}$ is minimally 2, which is the lowest possible sample size to compute a SD. The total number of animals used in then:

$$n_{total} = n_{exp} + n_{con}$$
$$n_{con} = n_{exp} - n_{prior}$$
$$n_{prior} = \sum_{p=1}^{p} n_p * index_p$$

where the number of animals of the control group ($n_{con}$) is diminished by the effective number of prior animals ($n_{prior}$), meaning the sum of the animals of each experiment used to define the prior ($n_p$) multiplied by the respective weight ($index_p$). The $index$ is a value between 0 and 1. An $index$ of 0.3 means that only 30% of the information in the prior study at hand will be used. In the simulation, we set $index = 1$. For analyses, researchers may opt to vary this value depending on the degree of similarity of the prior experiments to the current study. For more information about this topic, see *expert elicitation* (*12*).

In this simulation study, we assumed that the prior is a perfect estimation of the population. This issue will be further discussed in the "Sensitivity simulation" section.

**Validation study**

To validate the applicability of the method to real life situations (Fig. 2-B, Sub-section "RePAIR on RELACS"), *RePAIR* was applied to an experimental dataset (Sub-section "RELACS dataset") with a prior specified from unrelated literature (Sub-section "Prior specification").

RELACS dataset

To validate *RePAIR* on a real-life dataset, a well-powered dataset investigating a *real* and reproducible difference between two groups was required. We defined as "real" and "reproducible" an effect that persists in a high quality, well-powered meta-analysis. These criteria were met by the effects of early life adversity (ELA) on memory after non-stressful learning, as identified by a recent meta-analysis of literature previously conducted by our own lab (*5*). From this study, an effect size of *Hedge's G = 0.4* was estimated to describe the difference in performance on the memory task object in location between controls and animals that experienced ELA with the limited bedding and nesting (LBN) model (*16*). Considering a Welch two-sided independent means t-test and an alpha of 0.05, 200 animals would be required to achieve a power of 80%.

Due to the paucity of power of preclinical studies, it is not surprising that we were unable to identify any study on this experimental outcome using (at least) 200 animals. Even though no single laboratory works with such sample sizes, the required power could be reached by combining data of multiple laboratories. To this end, we created RELACS (*Rodent Early Life Adversity Consortium on Stress*), a unique rodent consortium constituted by several laboratories around the globe working on ELA. Since the scope of RELACS is broader than that of this study, we further specified inclusion criteria (Table S3) for the gathered data to be considered (and assumed) part of the same experiment. The criteria were specified *a priori* and blindly to the outcomes and the laboratories.

Within the RELACS consortium, 7 independent experiments met the specified criteria.
When analyzed independently, the p-value < 0.05 was reached in only 2 of the 7 experiments, in agreement with the low power of preclinical studies. By combining the 7 experiments, we reached a sample size of 275 animals, distributed as $n_{con} = 132$ and $n_{ELA} = 143$. The effect size *Hedge's G = 0.37* was similar to the one estimated from literature (*Hedge's G = 0.4*).

We concluded that this dataset meets the required criteria to validate *RePAIR*: it describes a reproducible effect as shown by the meta-analysis, and it is sufficiently powered since sample size is larger than the expected 200.

Prior specification

To mimic the profile of a junior scientist embarking on a new research project, one of us (VB) specified the prior to be used for the validation of *RePAIR* on the RELACS dataset. The researcher was asked to plan an experiment with the same characteristics (Table S3) as the RELACS dataset, i.e. investigating memory after non-stressful learning with the object in location task in adult male mice. The researcher was requested to select 8 publications that she would use to set up her study, while focusing on the control and not the experimental group. The selected publications did not belong to the ELA field, and were not used elsewhere in this manuscript. Furthermore, for each study the researcher defined a similarity index, a number between 0 and 1 that would express how similar the control group of each literature study was to the experiment that she was planning to perform (1 = identical). Two publications reported the same outcome on two separate groups of animals. Both experiments were considered, albeit with a lower index. The process was overseen by a senior researcher (RAS).

Although VB selected the prior blinded to the results of the RELACS dataset, it is important to notice that prior specification has some degree of subjectivity, i.e. another researcher may choose different publications on which to base their study (Section "Sensitivity simulation").

RePAIR on RELACS

By performing a Welch independent samples t-test (two-tailed, alpha = 0.05), we validated that the RELACS dataset supported that control and ELA groups differ in discrimination (p-value < 0.05). We performed several tests (Table S4) to confirm that by using *RePAIR* the same conclusion would be reached (Fig. 2-B).

Sensitivity simulation

The assumption that control groups come from the same population leads to the property that if sufficient samples (experiments) are available, the control population will be known. As the number and quality of experiments increases, the parameters identified will be better descriptions of the population. However, it is impossible to pre-define how many high-quality studies are necessary for an optimal parameters' estimation. Until optimal population parameters are known, prior specification may be subject to variation.

In our validation study, 10 experiments (from 8 studies) picked from literature (Section "Prior specification") had an extremely similar estimation of the population mean to the 7 experiments of the RELACS dataset (Fig S4-A). One may conclude that 7-to-10 experiments may be sufficient to estimate the population of our outcome. To test this hypothesis and to experimentally quantify relevant variation in article selection, we simulated many different priors by picking at random 10000 times $k$ experiments ($k$ equal to 2 through 16) from the 17 identified (of the validation study, 10 from "Prior" literature + 7 of RELACS dataset, Fig. S4-B). Variation in article selection for each $k$ was calculated as the 2.5$^{th}$ and 97.5$^{th}$ percentiles to avoid extreme values. Changes in *Hedge's G* between 0.1 and 0.5 could appropriately describe the variation across $k$ (Fig. S4-B), and 10 articles are here sufficient for a stable estimation of the population parameters. Of note, the sampling occurs from a finite population, where 17 experiments represent the reference value of the estimated variations. As a consequence, the intervals may be underestimated.

With this experimentally-derived estimation of population mean's variation, we conducted a sensitivity simulation study to investigate how prior control population mean's variation affects prospective study power (Fig. S4-D). Of note, this variation can act both in favor or against the hypothesis experimentally investigated (Fig. S4-C), depending on whether the prior control population mean moves towards or away from the population mean of the experimental group. Despite this limitation, we preferred this approach of experimentally deriving variation values over using a canonical variation of *Hedge's G = 0.1*

With the sampling strategy (Fig. S4-B), we sampled experiments, but, to keep consistency with the simulation study (Section "Using prior information"), the sensitivity simulation was conducted with number of animals. The relationship between number of experiments and number of animals is not straightforward. For example, one can achieve an $n_{prior} = 20$ with just one experiment, or two (e.g., each of $n = 10$), or three (e.g. $n = 9 + n = 6 + n = 5$). To transform variations due to experiments selection to variations linked to sample sizes, we identified across the $k$ * 10000 sampled estimations of means, animals roughly equivalent to 20, 50, 100, 200 ($n_{prior}$ in our sensitivity simulation (Section "Sensitivity simulation")). In these subgroups, we calculated the 2.5$^{th}$ and 97.5$^{th}$ percentiles, and visually validated their consistency with the results of Fig. S4-B. These values were used in the sensitivity simulation study to vary prior control population means (valued between 0 and ±0.5 *Hedge's G* depending on $n_{prior}$). All factors of the sensitivity simulation study were kept identical to the simulation study (Table S2) and here presented in the specific case of equal variances and effect size of *Hedge's G = 0.5.* The changes in prospective power were quantified and interpreted as a proxy of reliability of the method (Fig. S4-D). Since the estimated prospective power is consistently above the current median prospective power, *RePAIR* performs better than current practice, even in the worst (and unlikely) circumstances.

**Immediate potential impact**

By assuming that prior and study data come from the same population, we can reach more precise estimates of the parameters of the control population. This property can be used independently of how many animals are used and how well-powered an experiment is. Therefore, *RePAIR* can be applied immediately to animal experiments without necessarily increasing sample sizes, although this would probably be required and advisable.

To evaluate the immediate potential impact, we estimated the increase in prospective power if *RePAIR* would be used in new animal experiments with the resources currently available. We considered each study identified within each meta-analysis (Fig. S1 "Data A") as a new experiment where $n_{tot}$ was kept the same, but animals were redistributed in favor of the experimental group ($n_{exp} = 2 * n_{con}$, according to our rule of thumb). The controls of all other studies within the same meta-analysis were then considered as priors. In other words, $n_{prior}$ was calculated from the cumulative $n_{con}$ of all other papers included within the same meta-analysis. This cumulative $n_{prior}$ was then multiplied by the similarity $index = 0.3$, meaning that we valued the degree of similarity of the control groups of studies included in the meta-analysis to be 30%, and therefore we used only 30% of the information they hold. In this circumstance, the value of 0.3 is arbitrary. To evaluate how the similarity index affects power, we also calculated prospective power with a similarity index of 1 (Fig. S5). Prospective power was calculated (Fig 2-C) in the case of a Welch independent means t-test, for the plausible range of effect sizes previously identified (Fig. S2), when considering an alpha = 0.05. Since we adopted the same methodology and the same data (Sub-section "Prospective Power"), *RePAIR*'s immediate potential impact can be assessed by comparing the prospective power of Figure 1-C (without *RePAIR*) and Figure 2-C (with *RePAIR*). Briefly, *RePAIR* can substantially increase prospective power (i.g. from 12.5% to 68.7% for large effect sizes) even without increasing the total number of animals used, thereby directly addressing the insufficient statistical power of preclinical studies.

**Software used and created**

Every effort was made to minimize bias, e.g. data gathering and analysis was performed blindly, and multiple experts were consulted for sensitive information (inclusion/exclusion criteria). For data, R scripts and other information about the project, see https://osf.io/wvs7m/ . The following R packages were core to the project: 1) *tidyverse* for general data handling, 2) *shiny* for RePAIR webapp, 3) *MESS* for power calculations.

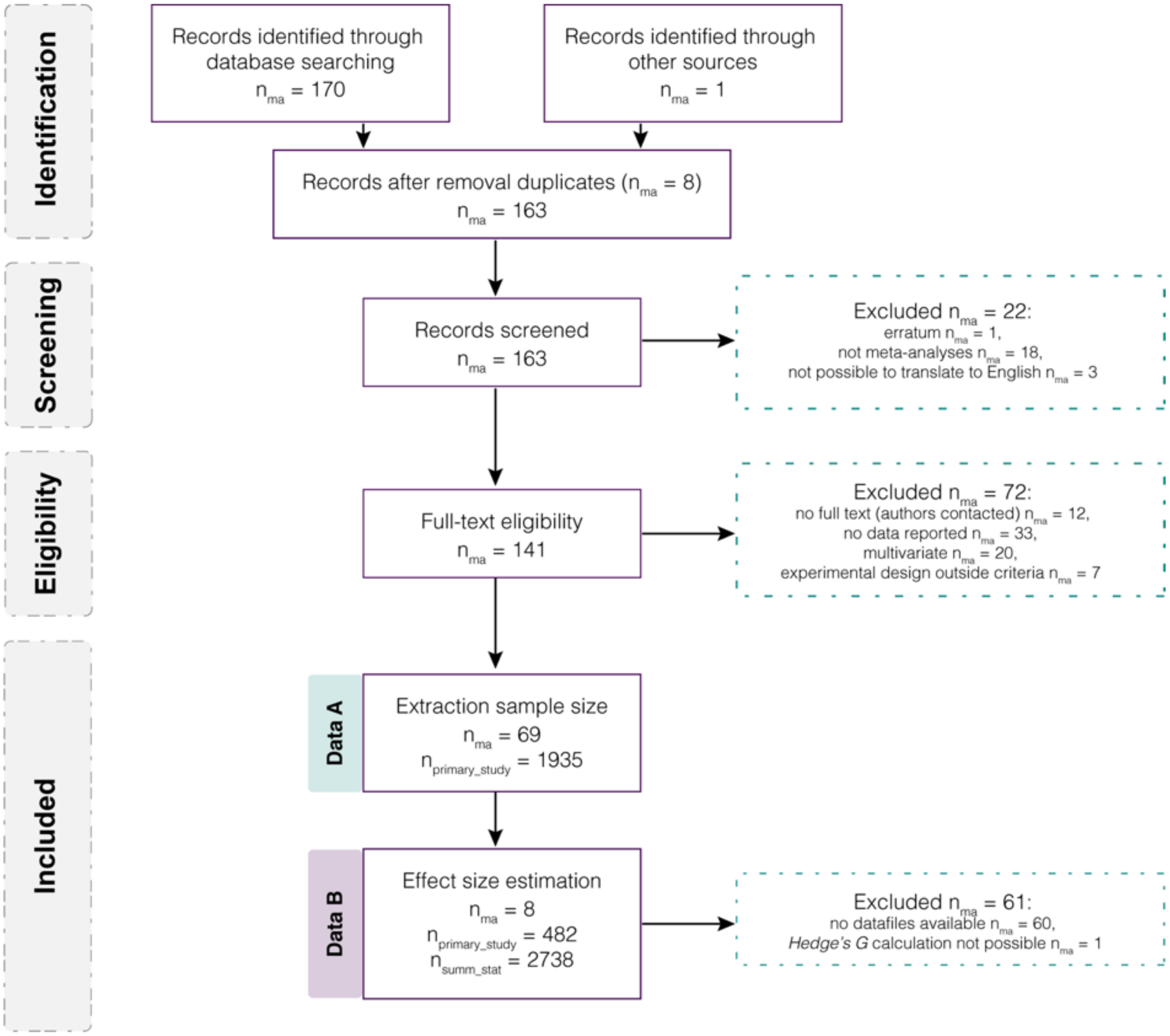

**Fig. S1.**

Flowchart methodology for data collection. From the EMBASE systematic literature search, 69 meta-analyses met our pre-specified inclusion criteria (Table S1). From the 1935 articles used in these meta-analyses, we extracted the sample size of the two largest groups ("DataA"), which was used for theoretical power calculations. 8 meta-analyses met our additional inclusion criteria (Table S1, "additional inclusion criteria"). From the 482 primary publications used in these meta-analyses, we extracted the available effect sizes, for a total of 2738 ("DataB"), which were used for calculations of post-hoc power (Fig.1-A) and range of effect sizes (Fig.S3). $n_{ma}$ = number of meta-analyses; $n_{primary\_study}$ = number of unique publications in mice and rats; $n_{summ\_stat}$ = number of summary statistics (mean, SD or SEM, sample size) extracted; Data A = data extracted at this level from dataset A; Data B = data extracted at this level from dataset B.

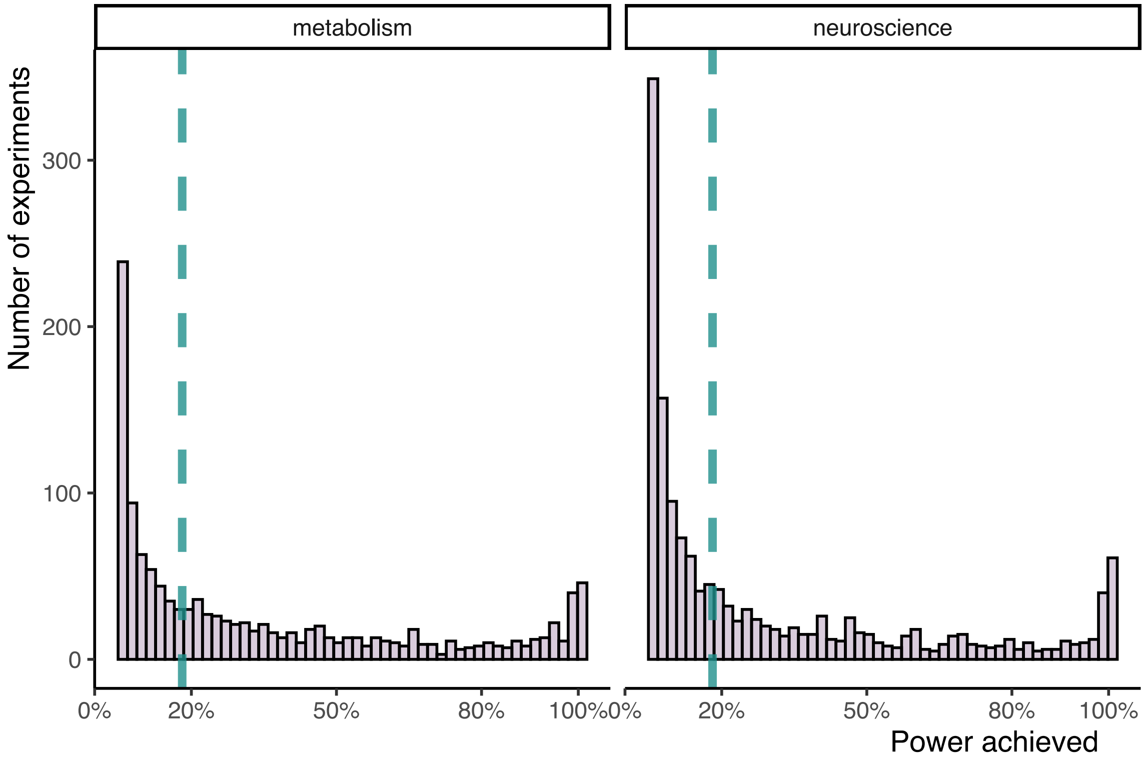

**Fig. S2.**
Median power achieved ("posthoc power") of rodent studies in Neuroscience and Metabolism research (Fig.S1 "DataB"). Green line: median of achieved power.

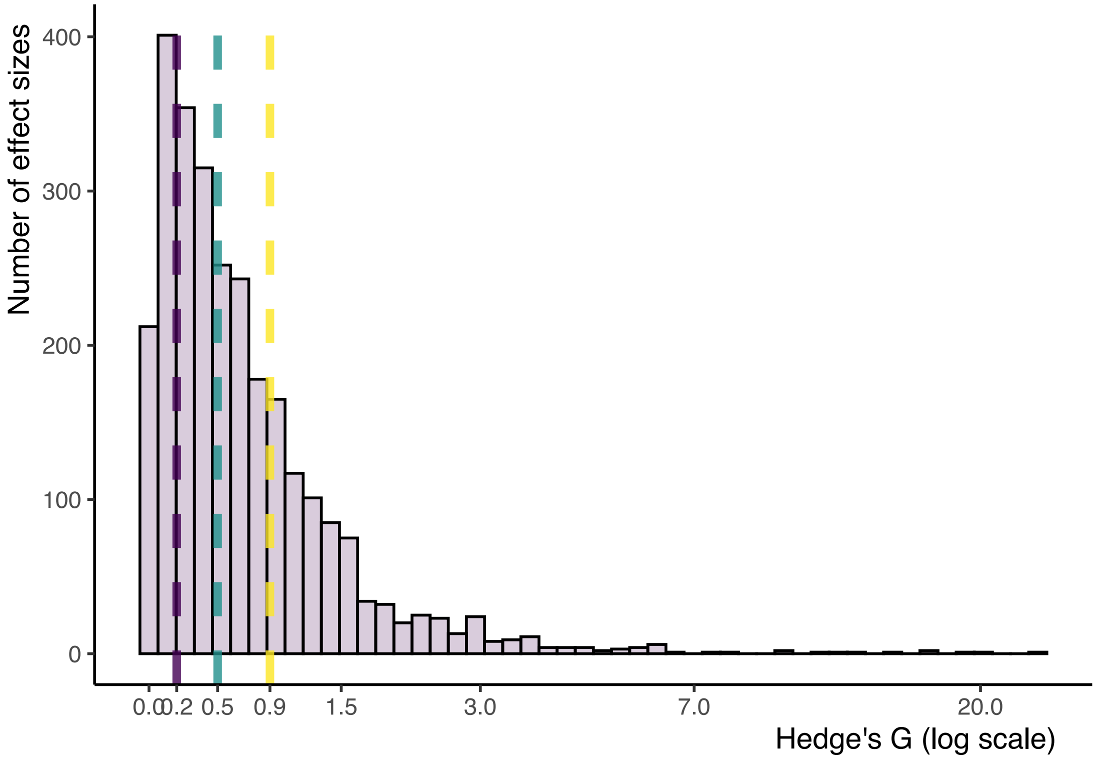

**Fig. S3.**

Range of common effect sizes estimated in preclinical literature. Frequency plot of experiments with a certain effect size Hedge's G. Purple line: 25th percentile; green line = 50th percentile (median); yellow line: 75th percentile. The related quantities (Hedge's G of 0.2, 0.5, 0.9) were defined as "small", "medium" and "large" effect sizes respectively. The graph is based on "Data B" (Fig. S1), but similar percentiles were replicated in a separate dataset (see R script, (*2, 10*)).

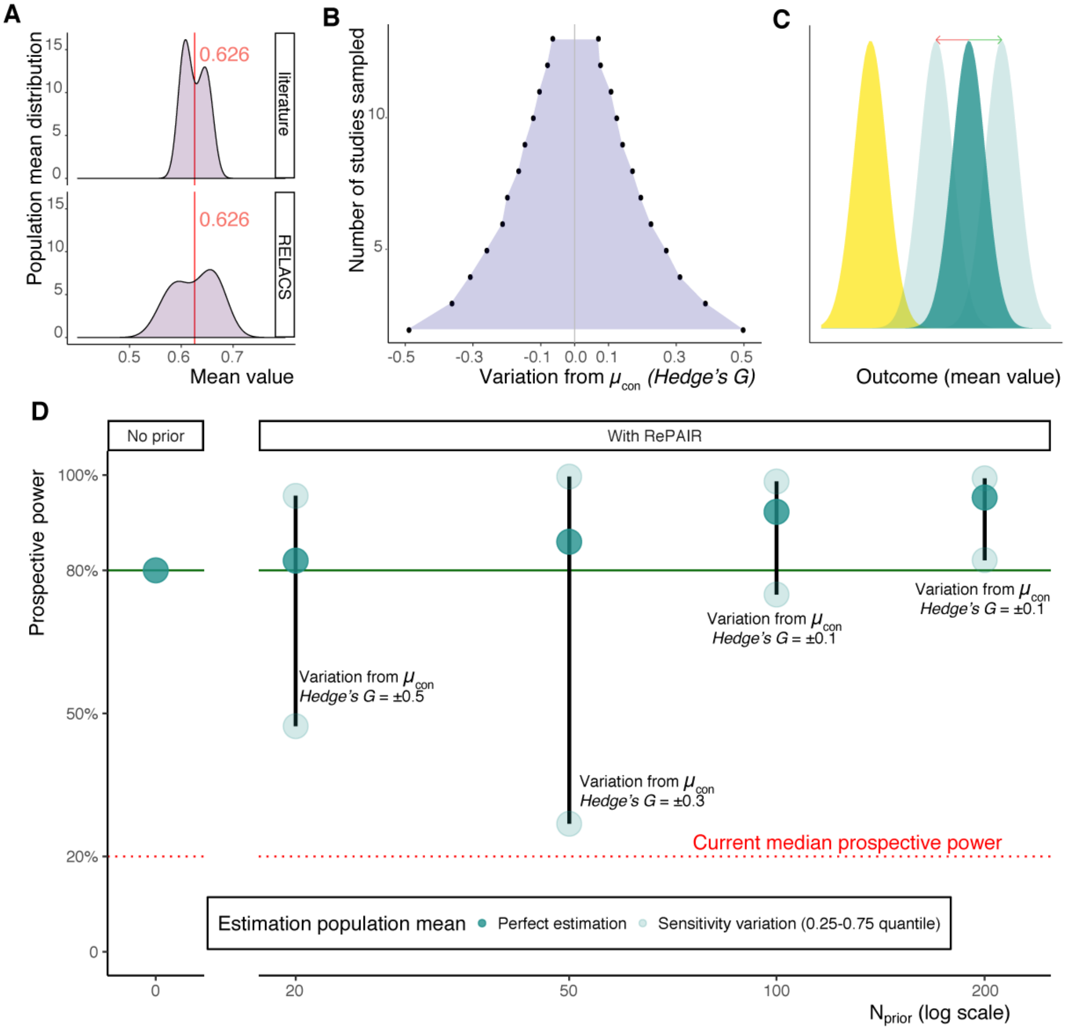

**Fig. S4.**

Sensitivity simulation. (**A**) Density distribution of control population means with data from literature (top, data from sub-section "Prior specification") and from RELACS (bottom). Red = mean of the control means. (**B**) Range of variation of estimation of populations means ($\mu_{con}$). To identify which deviations from the mean are relevant, we calculated the 2.5$^{th}$ and 97.5 percentile interval (grey area) of estimated means by sampling 10000 times 2 to 16 experiments from 17 studies (10 from literature (sub-section "Prior specification) and 7 from RELACS combined (A)). Once more than 10 experiments are used, the variation (*Hedge's G = 0.1*) becomes negligible. (**C**) Schematic representation of how variation of estimated population means can be both in favor (green arrow) and against (red arrow) the hypothesis scientifically investigated. Yellow = distribution of the experimental group; green = distribution of the control group. (**D**) Prospective power (*Hedge's G* = 0.5, equal variances) with RePAIR is higher than current practice (red dotted line) despite variations in population mean estimation (dots). Since $n_{tot}$ is not consistent due to the increasing of $n_{prior}$, prospective power can be interpreted only vertically (black line). Each vertical line displays how prospective power changes if the estimated prior mean is a perfect estimation of the population (dark dot), or deviates from it in favor (top light dot) or against (bottom light dot) the investigated hypothesis. The light dots correspond to rounded valued of the 2.5$^{th}$ and 97.5$^{th}$ interval calculated from (B) as specified in the sub-section "Sensitivity simulation". The exact variation for each percentile interval is written in figure. The progression of the dark green dots is an alternative visualization of the increase in green color intensity of Fig. 2-A.

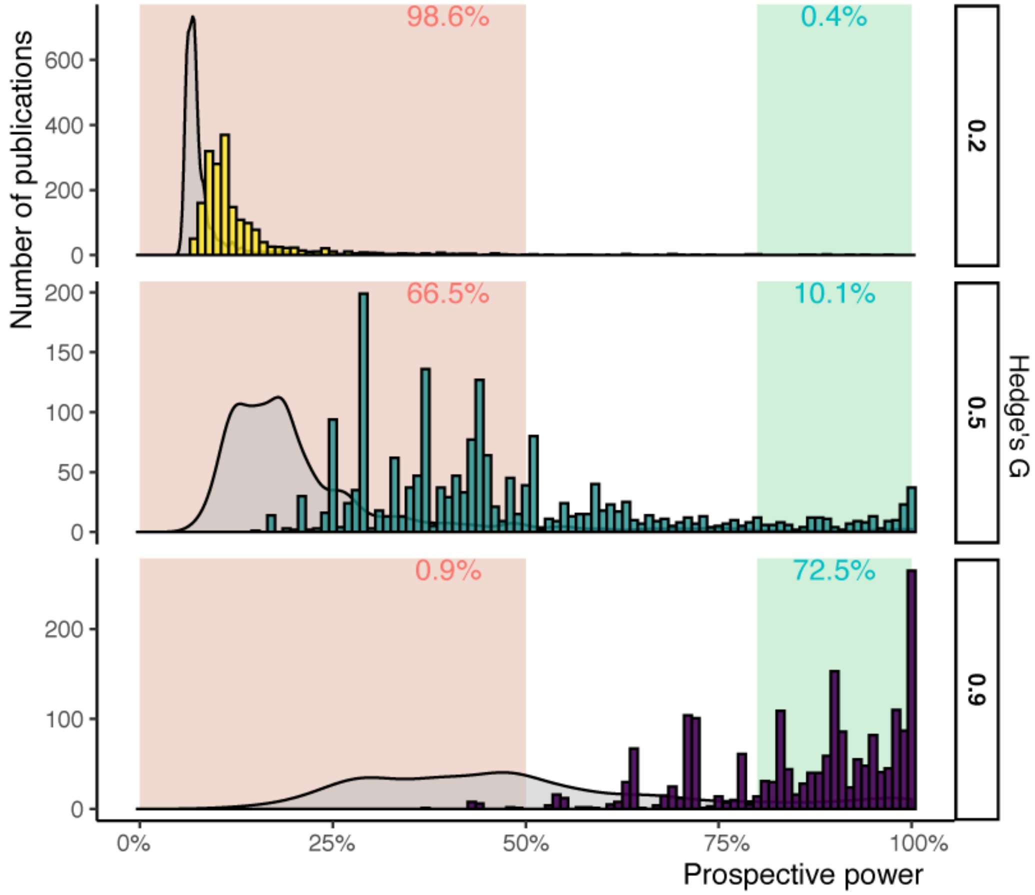

**Fig. S5.**
Prospective power of studies when RePAIR is applied to current resources redistributed in favor of the experimental group. Similarity index of 1. Graph based on “Data A” (Fig S1).

**Table S1.**

Inclusion and exclusion criteria for systematic literature search to identify relevant meta-analyses perform on data from mice and rats.

| **Inclusion** |
|---|
| Meta-analyses of literature |
| Mice and rats as population investigated |
| Comparison between (at least) two independent groups |

| **Additional inclusion criteria** | Only for range effect size estimation Fig. S2 |
|---|---|
| Data available as data file | |
| *Hedge's G* can be calculated with the available information | |

| **Exclusion** |
|---|
| Language not English or not translatable with google translate |
| Multivariate analyses (e.g. gene expression) |

**Table S2.**
Factors varied in the simulation study.

| Factors | Values and interpretation | Comments |
|---|---|---|
| $n_{prior}$ | 0 = noninformative<br>10 = pilot from own lab<br>20 = experiment from own lab<br>50 = routinely performed in own lab<br>100 = from literature or with data from other labs<br>200 = common outcome across labs | Values arbitrarily selected to simulate real-life situations |
| Effect sizes | *Hedge's G*<br>0.2 = small<br>0.5 = medium<br>0.9 = large | See Fig. S3 |
| Control population parameters | Standardized values:<br>$\mu_{con} = 0$<br>$\sigma^2_{con} = 1$ | |
| Experimental population parameters | **For mean:**<br>Calculated from *Hedge's G* definition<br>$$Hedge's\ G = \frac{\mu_{exp} - \mu_{con}}{\sigma^*_{pooled}}$$<br>Where<br>$$\sigma^*_{pooled} = \frac{\sigma^2_{exp} + \sigma^2_{con}}{2}$$<br>Therefore,<br>$\mu_{exp} = \mu_{con} + (Hedge's\ G * \sigma^*_{pooled})$ | Of note, μ and $\sigma^2$ to the population parameters. |
| | **For variance:**<br>Same variance as control, $\sigma^2_{exp} = \sigma^2_{con}$<br>Larger variance than control, $\sigma^2_{exp} = 2 * \sigma^2_{con}$ | For results of larger variance, see R script. [1] |
| Index | $index = 1$ | Not varied.[2] |

[1] Experimental groups are often more variables than controls; therefore, we performed the simulation also with unequal variances (equal variances are not assumed). Since the results of same and larger variances are extremely comparable, only same variances are reported in text. For larger variances, please see the R script.
[2] In the simulation, the *index* was not varied as it would simply decrease $n_{prior}$. For example, $n_{prior} = 50$ is equivalent to $n_{prior} = 100$ with an *index* = 0.5.

**Table S3.**
Inclusion/exclusion criteria for selection for RELACS dataset.

| Inclusion | Comments |
|---|---|
| Male mice | Insufficient data on females, males more frequently used |
| Adult | Older than 8 weeks of age but younger than 1 year |
| LBN as ELA model | Amount of bedding material of ELA animals could be both ¼ and ½ of controls. |
| Object in location task performed, with available exploratory time of both objects | Necessary to operationalize memory of each animal as: $discrimination = \frac{time_{novel}}{time_{old} + time_{novel}}$ Where $time$ refers to the time spent exploring an object in either the novel or old location. |
| Animals habituated to test cage prior to the learning phase | To avoid novelty-induced stress effects on memory |
| Exploration time of each object > 0s | Exploration of both objects is present |
| Preference of objects/locations avoided | Objects and locations were experimentally balanced or no preference was observed in previous experiments |
| Retention time between learning and test phase at least 1 hour | Working memory excluded |
| Experiments were performed and analyzed blindly and randomly | To ensure good experimental quality |

| Exclusion | |
|---|---|
| Metal grid not used in the LBN model | |
| Animals not habituated to test cage prior to the learning phase | To avoid novelty-induced stress effects on memory |
| Control group is unable to discriminate the novel location | Discrimination index unequal to 50% at a group level to exclude possible problems in the set-up of the experiment |

**Table S4.**

Summary of analyses to verify performance of *RePAIR* on RELACS dataset. This summary should be interpreted in relation to the results shown in Fig. 2-B.

| Fig.2-B | Aim | Experiment | Prior |
|---|---|---|---|
| *t-test* | RePAIR can achieve the same conclusion as a Welch t-test | RELACS | Non informative |
| *underpower* | When n_con is decreased, the test is no longer significant | ELA group from RELACS; control group 30% of RELACS (randomly selected) | |
| *Literature Prior RePAIR* | Prior of literature can substitute prior from same dataset | Same as *underpower* | From literature as selected by VB |

**List contributors to the RELACS consortium**

[&] collaborating authors to Rodent Early Life Adversity Consortium on Stress (alphabetical order):

M. Abbinck[5], T.Z. Baram[6], J.L. Bolton[6], J. Bordes[7], M. Joëls[1,4], J. Knop[1], A. Korosi[5], H. Krugers[5], J.T. Li[8], E. Naninck[5], K. Reemst[5], S.R. Ruigrok.[5], R.A. Sarabdjitsingh[1], M.V. Schmidt[7], E.H.L. Umeoka[5], C.D. Walker[9], X.D. Wang[10], K. Yam[5]

*[1] Department of Translational Neuroscience, University Medical Center Utrecht Brain Center, Utrecht University, The Netherlands.*
*[4] University of Groningen, University Medical Center Groningen, Groningen, The Netherlands.*
*[5] Swammerdam Institute for Life Sciences, SILS-CNS, University of Amsterdam, Amsterdam.*
*[6] Department of Anatomy/Neurobiology, University of California, Irvine, Irvine, California; Department of Pediatrics, University of California, Irvine, Irvine, California.*
*[7] Max Planck Institute of Psychiatry, Department of Stress Neurobiology and Neurogenetics, Munich.*
*[8] Department of Neurobiology, Key Laboratory of Medical Neurobiology of Ministry of Health of China, Zhejiang Province Key Laboratory of Neurobiology, Zhejiang University School of Medicine, Hangzhou, China.*
*[9] Dept of Psychiatry, McGill University, Douglas Institute Research Center, Montreal, Quebec, Canada.*
*[10] Clinical Psychopharmacology Division, National Clinical Research Center for Mental Disorders (Peking University Sixth Hospital/Institute of Mental Health) and the Key Laboratory of Mental Health, Ministry of Health (Peking University), Beijing, China.*
